# The role of tactile interactions in flight responses in the Bronze Cory catfish *(Corydoras aeneus)*

**DOI:** 10.1101/449272

**Authors:** RJ Riley, ER Gillie, A Jungwirth, J Savage, NJ Boogert, A Manica

## Abstract

One of the primary functions of animal aggregations is defense against predators. Many social animals enjoy reduced predation risk as a result of grouping, and individuals within groups can benefit from information transferred by their group-mates about a potential predator. We present evidence that a tactile interaction behavior we term ‘nudging’ substantially modified group responses to a potential threat in a highly social catfish species, *Corydoras aeneus*. These catfish deployed nudges during flight responses, and these nudges were associated with a greater likelihood of group cohesion following a threat event. Increased nudging behavior resulted in longer flight responses. In addition, individuals that perceived the threat first were more likely to initiate nudges, implying that nudges could be used to alert group-mates to the presence of a threat. Taken together, our results suggest that tactile communication plays an important role in gaining anti-predator benefits from sociality in these fish.

## Introduction

Animal aggregations occur across the animal kingdom, with this ubiquitous sociality likely arising through the profound advantages group living can offer. Among the most evident of these benefits is reducing the risk of predation (Neill and Cullen, 1974, Major 1978). Aggregative behaviors that reduce predation risk are observed in invertebrates such as aphids (Turchin, 1989), in many fishes including minnows (Pitcher, 1986) and guppies (Magurran, 1994), in reptiles such as iguanas (Greene, 1978), in many birds including cliff swallows (Brown, 1988) and ostriches (Bertram, 1980), and in many mammals including prairie dogs (Hoogland, 1981). Predation risk also increases cohesion in many species, including walleye (Sogard, 1997) and fiddler crabs in the context of a ‘selfish herd’ response (Viscido, 2002). By living in close proximity to others, individuals gain the benefit of their conspecifics’ perception and attention, and can dedicate less time to predator vigilance while still being more likely to escape an attack (Bertram, 1980, Hoogland, 1981).

In addition to the benefit of seeing through their group-mates’ eyes, individuals also avoid predators due to the active spread of information about potential threats through groups. In some species, explicit signals such as alarm calls are delivered by one individual to alert its conspecifics about a predator, as in primates such as vervet monkeys (Seyfarth, 1980) and birds such as black-capped chickadees (Templeton, 2005); in the prior two examples, alarm calls encode specific information about the predator that is utilized by the caller’s conspecifics. In other systems, including fathead minnows and zebrafish, injured individuals release an alarm pheromone that alerts conspecifics to danger, albeit without specific information about the predator (Stensmyr, 2012, Brown, 2001). Predator inspection is another behavior that occurs in many taxa, including birds (Hinde, 1954) and fishes (Pitcher, 1986), and is performed by individuals or sub-groups but has consequences for the entire group. For example, in minnows a small contingent of a much larger shoal will inspect a potential predator at great potential risk, and if they perceive that the predator is a threat, they return to the shoal instead of hiding immediately, after which their shoal-mates alter their behavior (Pitcher, 1986). This implies that information about the predator was transferred, although it seems the individuals rely mostly on personal information to ignite a flight response (Magurran, 1988).

The acquisition of information from conspecifics that is potentially costly to obtain personally is certainly beneficial to individuals, and many species have evolved social behaviors that allow individuals to convey information about predators to their group-mates or otherwise influence their group-mates’ behavior in mutually beneficial ways. Understanding how an individual’s behavior can impact the coordination of its group is of particular importance to understanding how groups function, and how group living provides the myriad of advantages we see across taxa. One factor that strongly affects group coordination is familiarity, defined as previous experience with a given other individual. Familiarity leads to improved coordination in a variety of taxa, including birds (Senar, 1990) and schooling fishes (Ward, 2003). In particular, familiarity improves a group’s anti-predator defenses, such as in great tits, in which previous experience with nest-site neighbors results in a higher probability of a familiar neighbor contributing to the defense of a conspecific’s nest (Grabowska-Zhang, 2012). In fathead minnows, cohesion is greater and anti-predator behaviors (i.e. predator inspection) more effective in familiar groups when compared to unfamiliar groups (Chivers, 1995), and the same effect has been observed in juvenile trout (Griffiths, 2004). Given the advantages of familiarity, it makes sense that individuals often prefer familiar individuals over unfamiliar ones in a number of species, including cowbirds (Kohn, 2015) and guppies (Griffiths, 1997).

This study investigated how individual *Corydoras aeneus*, the Bronze Cory catfish, can initiate or mediate a coordinated group response to a potential predator attack, how familiarity affects interactions during a group response to a predator attack, and how individuals can maximize group coordination under stressful circumstances. The Bronze Cory catfish is a highly social neotropical species (Lambourne, 1995) that uses an unusual tactile interaction behavior in which individuals physically nudge one another during coordinated movements (Riley, 2018). To investigate how individuals use nudges in response to a potential threat, and how nudges affect group coordination, we scrutinized this behavior in a controlled laboratory setting. We predict that familiarity will impact nudging tendencies, and individuals will be more likely to both deliver nudges and successfully recruit their familiar partner over their unfamiliar partner. We also predict that nudging may serve an important function for both the spread of information about a potential predator, as well as maintaining proximity between group members following a flight response.

## Methods

### Study species

In the wild, *Corydoras aeneus* are social foragers that live in groups of variable size consisting of males, females, and juveniles (Lambourne, 1995). Because they are bottom-dwelling they shoal in 2 dimensions and their social behaviour can be accurately and reliably recorded from above. We have observed that both wild-caught and captive-bred individuals exhibit an unusual tactile interaction behaviour during coordinated activities and following a startle response in aquarium settings (as in Riley, 2018). Wild fish were observed utilizing this behavior in several small streams in the Madre de Dios locality of the Peruvian Amazon in 2011 and 2013, particularly when fleeing the observer (RJR, pers. obs.).

### Husbandry

We obtained Bronze Cory catfish from three local pet shops in Cambridgeshire: Maidenhead Aquatics Cambridge, Pet Paks LTD, and Ely Aquatics and Reptiles. All fish used in both experiments were at least 24 weeks of age, and had been housed in the lab for at least six weeks prior to the start of experiments. We maintained the fish on reverse osmosis (RO) water purified to 15ppm or less total dissolved solids (TDS) and re-mineralized to 105-110ppm TDS using a commercially prepared RO re-mineralizing mix (Tropic Marin Re-mineral Tropic). The fish lived on a 12:12 light:dark cycle at a temperature of 23 +/− 1 degree Celsius. Prior to the start of the experiment, we housed the fish in mixed-sex social housing tanks (60cmx30cmx34cm) of 6-10 fish. The tanks were equipped with 4 Interpet Mini internal filters and an air stone. We fed the fish daily on a varied diet of alternating Hikari wafers (Hikari brand, USA), Tetra Prima granules (Tetra brand, Germany), and thawed frozen bloodworms (SuperFish, UK). The group composition of social housing tanks was stable for at least six weeks prior to experiment, and unfamiliar fish (see below) had not been exposed to each other for at least six months prior to the experiment, if at all. At the conclusion of each experiment, all fish were returned to their respective social housing tanks.

### Triplet study experiment procedure

We investigated the behavior of triplets over three weeks in May-June 2017, as in (Riley, 2018). We analyzed 18 triplets for a total of 54 individuals. Each triplet consisted of two familiar individuals taken from the same social housing tank and an unfamiliar individual taken from a different tank. Triplets were composed of same-sex individuals to avoid courtship interactions, and fish were not fed prior to the trial to encourage exploratory movement in search of food. Each triplet was placed in a testing arena situated on a very low shelf 3cm off of the floor (Figure 1; two identical setups were utilized in parallel). The arena had water 29 cm deep and a thin layer of aquarium sand as substrate (mimicking the Bronze Cory catfish’s natural habitat) and an opaque barrier so that each open (initially partitioned) arena was (47cm x 30 cm). We placed a piece of opaque acrylic outside the half of the tank where a sheltered ‘cover’ area was provided so that fish could not see any potentially threatening stimuli from outside the tank while in cover. Fish were left in the open part of the arena for an hour to acclimate to their group-mates, and were filmed during this time so that nudging patterns at ‘baseline’ (in the absence of threat stimuli) could be analyzed. The partition was then removed, and fish were allowed to explore the entire tank (91cm x 30cm) for 30 minutes prior to the threat event period in order to explore the testing arena and become familiar with the location of cover. We used a GOPRO HERO 3 camera to film each triplet for the entire duration of the acclimation periods and threat events.

**Figure 1:**
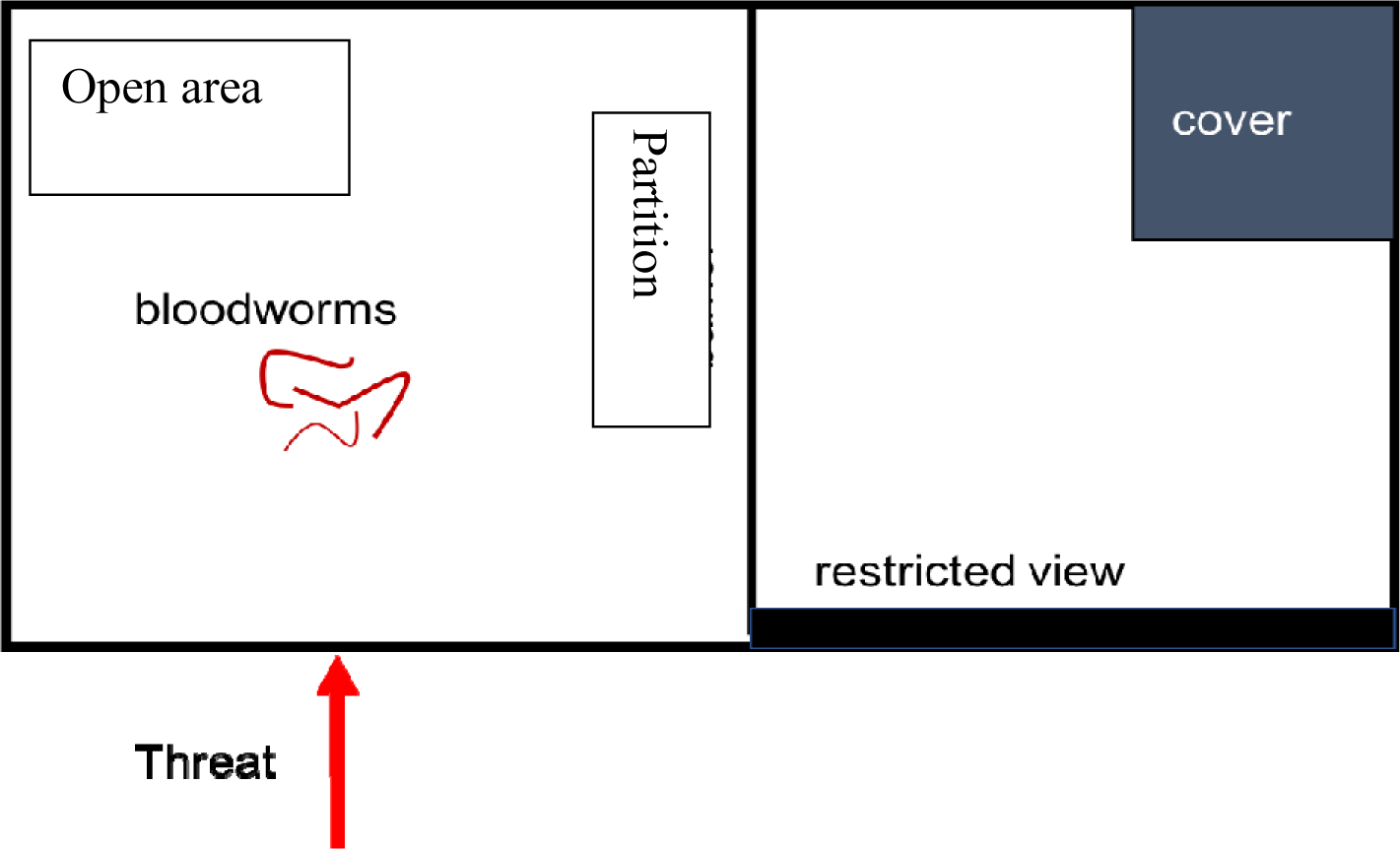
Schematic of test tank. Fish were initially allowed to acclimate in the open (initially partitioned) area without food. After acclimation, the barrier was removed, and fish were allowed to explore the entire test tank. Bloodworms were then added to the open area to encourage fish to leave cover. The threat stimulus was only applied when fish were in the open area.

### Providing the threat stimulus

After fish were allowed to explore the entire test tank, i.e. 1.5 hours after introduction, the threat event period commenced. Threat events were delivered by RJR or ERG and were given through rapid approach of the test tank from a distance of 1.5 meters. Threat events were delivered in similar clothing every day (blue jeans and the same shoes during each event) and happened with an approach speed of roughly 2-2.5 m/s. Triplets responded to the vast majority of these threat events with a clear anti-predator response, and most responses to threat events fell into two categories: (i) fish responded to the threat by swimming rapidly to cover, often at speeds that required frame-by-frame video analysis for data extraction. (ii) fish responded to the threat with rapid movement, then froze in place outside of cover. We considered both reaching cover and freezing (following an initial burst of movement) as complete threat responses, thus considering a given threat event as ‘complete’ once all fish of a triplet had fled to cover or frozen. Following each threat event, we used an aquarium net to chase into cover any individuals in the exposed area. If the aquarium net appeared before all individuals had frozen or reached cover, the threat was considered incomplete.

Threats were only delivered when all fish were out of cover in the open area. Thawed frozen bloodworms were delivered to the open area to provide an incentive for fish to leave cover. Additional bloodworms were provided after threat events. Triplets were allowed to recover in between threat events, which occurred at least 4 minutes apart. Groups varied in how long they spent in the open area, as well as how long they needed to recover from the threat event and resume normal activities following a threat event, and the threat event period lasted a maximum of 4.5 hours. After the threat event period, all individuals were returned to social housing tanks. Consequently, the total number of threats each triplet received differed.

### Analysis

### Recorded behaviors

For each triplet, we recorded (1) how many threat events took place, (2) the order that the group responded in (i.e. which individual responded first, second, or third), (3) how many times each group member nudged each of its groupmates, (4) whether any given nudge resulted in a previously stationary fish initiating a flight response, a case we defined as ‘recruitment’, and (5) the ‘flight time’ of each individual, defined as the time taken by each individual to either reach cover or freeze; ‘flight time’ refers to the duration of each individual’s flight response, starting at the onset of the threat event and ending when the individual reached cover or froze in place. For our analysis of mean flight times, we only used threat events that were considered complete. Finally, we assessed general measures of cohesion. These include whether or not all three group members were in proximity (within 7cm, or roughly two body lengths) to one another 30 seconds before the threat, three seconds before the threat, and three seconds after the threat event.

### Analysis

All statistical analyses were carried out in R version 3.2.2 (R core developer team), and generalized linear mixed effects models (GLMMs) were fitted using the lme4 package (Bates et al 2013). All GLMMs were used to investigate count data and were thus fitted assuming a Poisson error distribution.

To test whether a triplet’s baseline nudging tendency was correlated with their nudging tendency during flight responses, we used a Spearman’s rank correlation test. We tested the correlation between a triplet’s total number of nudges during the acclimation phase and that triplet’s average number of nudges per threat event.

We used a GLMM to analyse whether familiarity influenced individual nudging preferences during threat events. The model included the number of nudges an individual initiated as response variable, the familiarity between initiator and receiver as explanatory variable (binary: receiver familiar or unfamiliar to the initiator), and two random effects (initiator ID and group ID).

Similarly, to analyse whether familiarity influenced recruitment rates during threat events, we used a GLMM with the number of successful recruitments by an individual as response variable. As above, the model included the familiarity between initiator and recruit as explanatory variables (binary: recruit familiar or unfamiliar to initiator), and two random effects (initiator ID and group ID).

To test whether nudging frequency during threat events was correlated with the probability of group cohesion following a threat event, we used a Spearman’s rank correlation test. We tested the correlation between the average number of nudges a triplet performed during threat events and the proportion of all threat responses that ended with that triplet showing group cohesion (as defined above).

We analysed whether nudging rate during threat events influenced flight times and/or whether nudging rates changed throughout consecutive exposures to threat events using a further GLMM. The model included median flight time during a threat event (i.e. the median time it took a triplet’s individuals to complete their threat response to a given threat event) as an explanatory variable, the sequence of the treat event as an explanatory variable, the total number of nudges performed by the triplet during the threat event as the response variable, and group ID as random effect.

Finally, to analyse whether the order in which individuals of a triplet reacted to the threat influenced the number of nudges a given individual initiated and/or whether this changed throughout consecutive exposures to threat events, we used a GLMM. The model included the total number of nudges an individual initiated during a given threat event as response variable, two explanatory variables (that individual’s rank in the order of response: 1^st^, 2^nd^, 3^rd^ responder; count of threat events the triplet had experienced), and group ID as random effect.

## Results

### Comparison to baseline

Group nudging behavior at baseline was significantly correlated with the average number of nudges per threat event (Spearman’s rank correlation, S = 427.44, p-value = 0.016, figure 2). Groups that nudged more during a set duration while exploring in the absence of simulated threats tended to nudge more during threat events.

**Figure 2:**
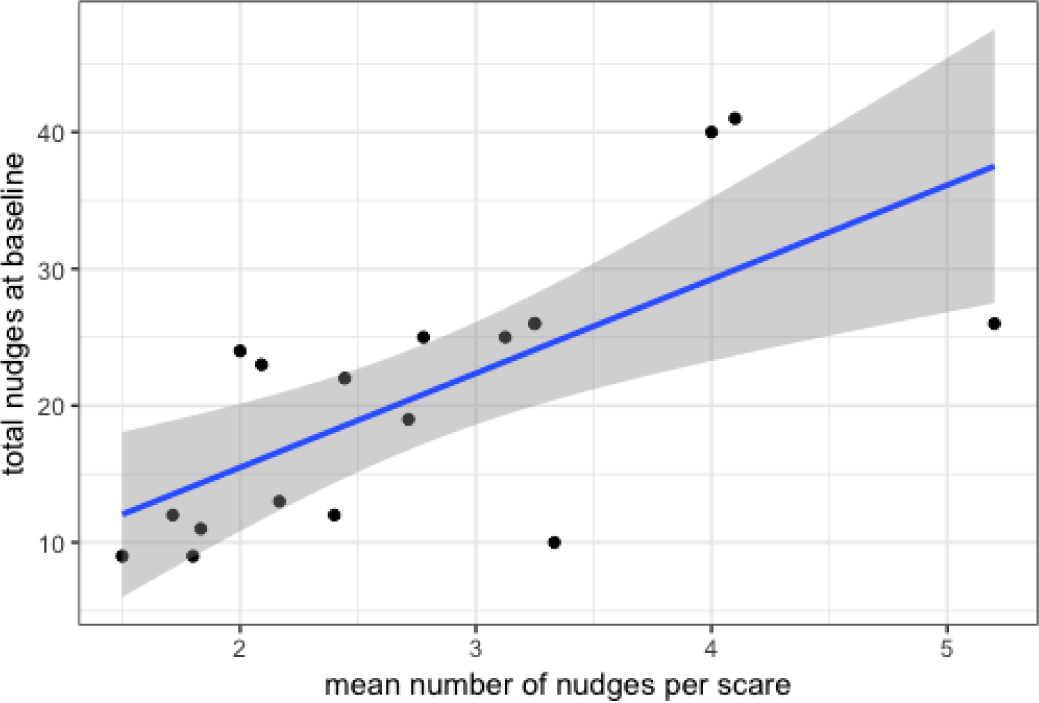
the mean number of nudges per threat event vs the total nudges at baseline. The blue line represents the line of best fit for this relationship. The shaded area represents the 95% confidence interval

### General

Overall, out of 135 threat events, 121 threat events were complete (all fish of a triplet fled to cover or froze in place). The 14 incomplete events involve fish either not responding to the stimulus at all, or still being in motion by the time the aquarium net chased them to cover. Out of the 121 complete threat events, 105 threat events involved one or more nudges. Individuals displayed no preference for delivering nudges with familiar partners as opposed to unfamiliar partners (Poisson generalized mixed effects model with individual and group ID as random effects, χ^2^_1_=2.8, p=0.093). There was also no effect of familiarity on nudge frequencies during baseline (Riley, 2018). The proportion of nudges that resulted in recruitment was 0.26. Individuals did not respond with higher recruitment rates to nudges received from familiar vs unfamiliar group-mates. (GLMM with individual and group ID as random effects, χ^2^_1_= 1.11, p=0.292)

We found a significant correlation between a group’s mean number of nudges per threat event and the proportion of threat events where all group-members were together three seconds following the event (Spearman’s rank correlation, S = 335.24, p-value = 0.003, Figure 3). Groups that had a higher mean number of nudges per threat event exhibited higher cohesion following the threat event.

**Figure 3:**
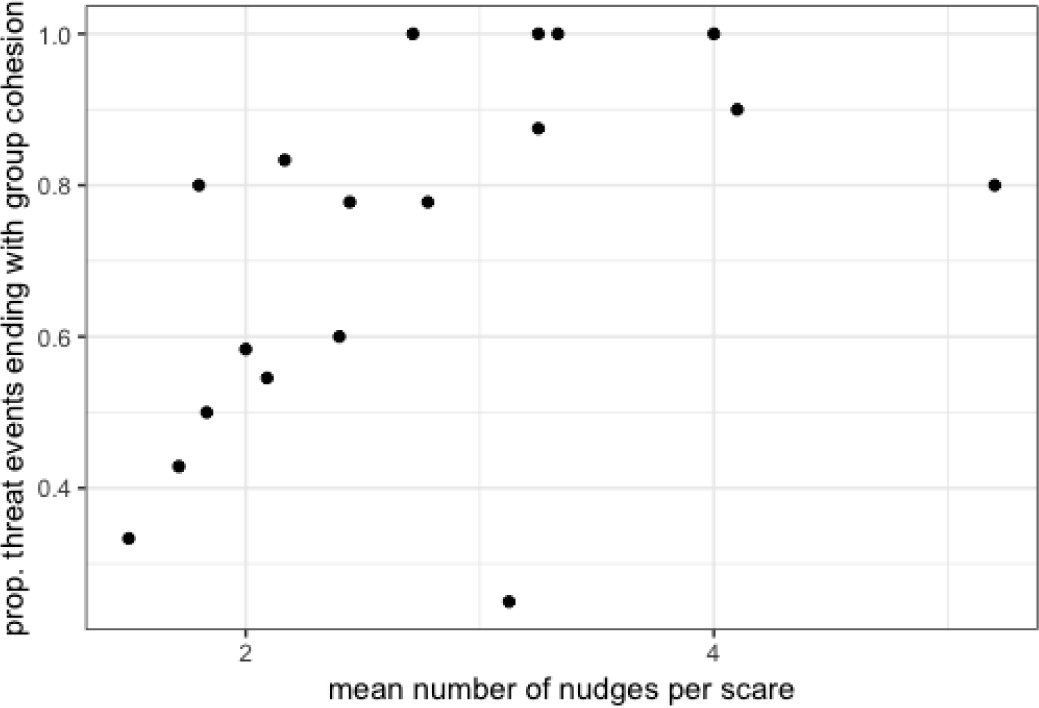
proportion of threats that ended with all group members together vs mean nudges per threat event

Threat events in which group members exhibited more nudges had longer flight times (generalized mixed-effects model with Poisson error structure; Group ID as a random effect, χ^2^_1_= 43.6, p<0.001, Figure 4.) The threat event number did not have a significant effect on group median flight time (χ^2^_1_=2.4, p=0.120)

**Figure 4:**
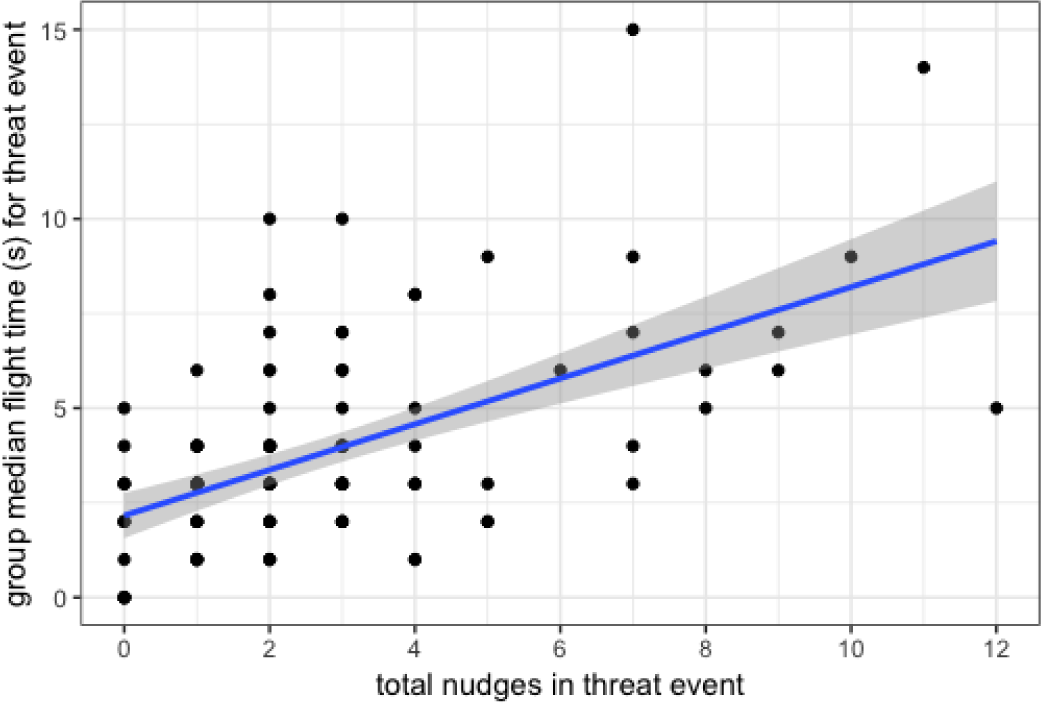
group median flight time total nudges in threat event. The blue line represents the line of best fit. The shaded region represents the 95% confidence interval

The sequence in which individuals responded to the threat event had an effect on the number of nudge initiations: earlier responders initiated more nudges (generalized mixed-effects model with Poisson-distributed error structure; Group ID as a random effect χ^2^_1_=4.6, p=0.032, figure 5. The scare event number did not have a significant effect on the number of nudges an individual initiated (LRT=2.86, df=1, p=0.09).

**Figure 5:**
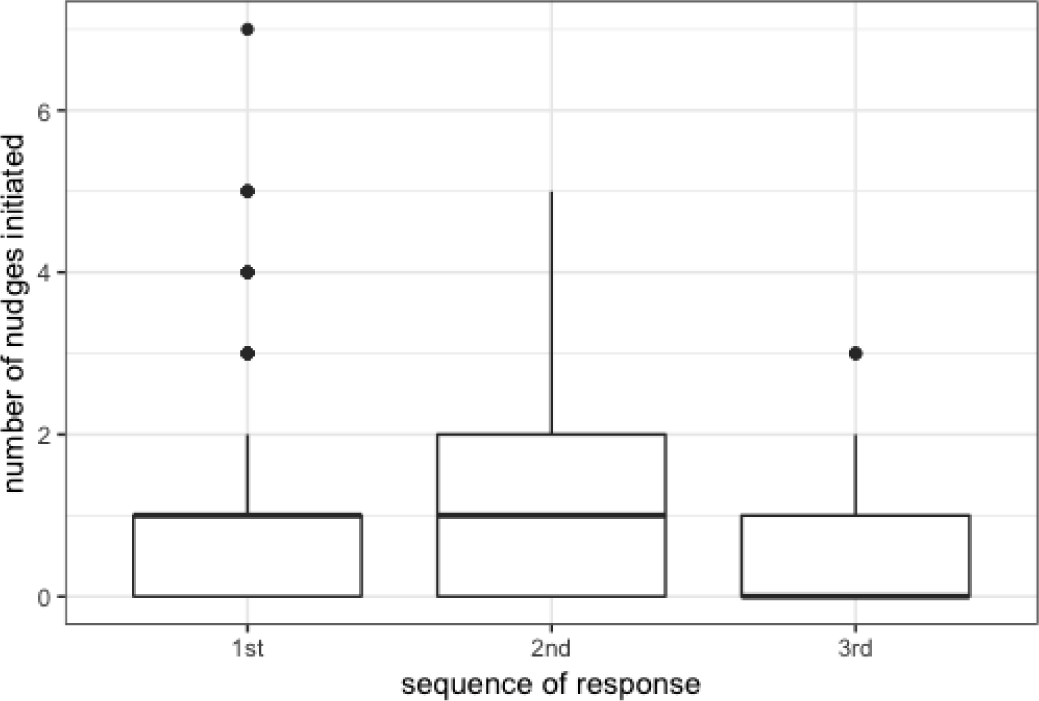
nudges initiated by the first, second, and third fish to respond to a threat.

## Discussion

Our results show that, for triplets, nudging patterns are consistent across context, and groups that nudge more frequently during environmental exploration nudge more frequently during a flight response. Familiarity of group-mates did not impact nudging behavior, and unfamiliar and familiar individuals were equally likely to initiate or receive nudges following a potential threat. This is somewhat surprising, as familiarity has clear effects on individual preference in other taxa, and individuals are more likely to assist the defense of familiar conspecifics (Grabowska-Zhang, 2012). However, given the high costs associated with a potential predator attack, and the fact that the Bronze Cory catfish have been shown to use nudging to compensate for lack of familiarity when foraging in the absence of a threat (Riley, 2018) it seems likely that the use of nudges by Bronze Cory catfish make familiarity irrelevant in such high-stakes circumstances.

In Cories, nudging was beneficial to all group members, and had a potentially selfish advantage for the initiator of the nudge. During a flight response, nudges were positively associated with a higher likelihood of maintaining cohesion following threat events and longer flight times. Meanwhile, an individual’s early detection of a threat relative to its group-mates is associated with initiating more nudges during the group’s threat response. These results suggest that individuals deploy nudging in response to potential threats, and that it fundamentally alters a group’s behavior by increasing the probability of group cohesion. For this reason, familiarity may not affect an individual’s decision of who to nudge, as the incentive to maintain cohesion is paramount. Furthermore, Bronze Cory catfish nudge group-mates extensively regardless of familiarity, and can also use increased nudging to coordinate effectively with unfamiliar partners (Riley, 2018). Given the serious consequences of a potential predator attack and the ubiquity of nudging directed to both familiar and unfamiliar individuals, it is perhaps practical that familiarity does not affect the flight response. This mirrors findings in other systems, such as rats, in which reciprocity of cooperative behaviors is related solely to prior experience of cooperation, and not familiarity with the current beneficiary of cooperation (Rutte, 2007).

Furthermore, social animals with an incentive to maintain proximity with conspecifics have often been shown to use behaviors to coordinate with others. In green woodhoopoes, vocalizations are used to maintain group cohesion while moving to new territory (Radford, 2004), and white-tailed deer exhibit a low-cost flagging alarm signal to recruit other individuals to join it in a flight response to a potential predator (LaGory, 1987). The Bronze Cory catfish also appears to utilize an interactive behavior in order to influence the dynamics of its group following a flight response to a potential threat. The nature of this behavior, a tactile nudge, perhaps lends itself to maintaining cohesion, as the initiator must be in such close physical proximity as to touch the receiver of its nudge.

Bronze Cory catfish may have evolved this behavior for a variety of reasons. They tend to live in small, murky streams (Nijssen in Lambourne, 1995), which we observed at our field site (RJR, personal observation), and were observed in other experiments to have poor vision (Kohda 2002), a characteristic we have also noted in wild and laboratory populations. Under these conditions, living in shallow water with poor visibility, a tactile mode of interacting with one another might be the most effective way for individuals to transfer information and maintain contact. This may also allow groups to maintain higher levels of cohesion: if an individual loses contact with its group-mates, it might be difficult to find them again, and individuals will be vulnerable while they search for one another. Tactile communication is present in another interspecific association, the well-documented shrimp-goby association, under similar circumstances, as the shrimp in that association has relatively poor vision (Kramer, 2009). In this system, shrimps convey their location outside the burrow by touching the goby with their antennae, and gobies, who have superior vision and serve as lookouts for predators, convey information about predators to their shrimp via a flick of the tail, a tactile signal that the shrimp can perceive, and after which both the shrimp and goby take shelter inside the burrow dug by the shrimp (Preston, 1978).

In this way, the Cories’ poor eyesight may have led to the evolution of an intraspecific tactile interaction method that can be deployed to spread information about predators and maintain cohesion following an attack. Individuals that reacted to a threat earlier initiated more nudges, which implies that this behavior may be used more frequently by individuals who have already perceived the threat and are altering their behavior in a way that transfers information to a group-mate. The fact that some of these nudges resulted in ‘recruitment’ in the sense that the receiver had been stationary prior to the nudge and then initiated a flight response following the nudge implies that nudges can alter the behavior of receivers and potentially alert them to the presence of a threat. It seems very likely that nudges modify the behavior of groups following a potential threat, and provide the benefit of increasing the likelihood of maintaining group cohesion while also encouraging a group to flee from a potentially dangerous area. Our study suggests that the Bronze Cory catfish’s nudging behavior can provide benefits when groups are responding to a potential attack. Nudges were also associated with coordinated movements in pairs of fish (Riley, 2018), and the consistency in nudging in triplets during exploration/foraging and while responding to potential threats suggests that nudging is a behavior that is useful for coordination under different conditions, and that nudging more frequently during exploration/foraging may contribute to a group’s coordination during responses to a potential threat. In this way, nudging appears to be a versatile behavior that plays a pivotal role in many aspects of the Bronze Cory catfish’s social life.

## References

Bergstrom, Carl T., and Michael Lachmann. 2001. “Alarm Calls as Costly Signals of Anti-Predator Vigilance: The Watchful Babbler Game.” https://digital.lib.washington.edu:443/researchworks/handle/1773/2000.

Bertram, Brian C. R. 1980. “Vigilance and Group Size in Ostriches.” Animal Behaviour 28 (1): 278–86. https://doi.org/10.1016/S0003-3472(80)80030-3.

Brown, Charles R. 1988. “Social Foraging in Cliff Swallows: Local Enhancement, Risk Sensitivity, Competition and the Avoidance of Predators.” Animal Behaviour 36 (3): 780–92. https://doi.org/10.1016/S0003-3472(88)80161-1.

Greene, Harry W., Gordon M. Burghardt, Beverly A. Dugan, and A. Stanley Rand. 1978. “Predation and the Defensive Behavior of Green Iguanas (Reptilia, Lacertilia, Iguanidae).” Journal of Herpetology 12 (2): 169–76. https://doi.org/10.2307/1563404.

Hinde, Robert Aubrey. 1954. “Factors Governing the Changes in Strength of a Partially Inborn Response, as Shown by the Mobbing Behaviour of the Chaffinch (Fringilla Coelehs) II. The Waning of the Response.” Proc. R. Soc. Lond. B 142 (908): 331–58. https://doi.org/10.1098/rspb.1954.0029.

Hoogland, John L. 1981. “The Evolution of Coloniality in White-Tailed and Black-Tailed Prairie Dogs (Sciuridae: Cynomys Leucurus and C. Ludovicianus).” Ecology 62 (1): 252–72. https://doi.org/10.2307/1936685.

Kohda, Masanori, Kanako Yonebayashi, Miyako Nakamura, Nobuhiro Ohnishi, Satoko Seki, Daisuke Takahashi, and Tomohiro Takeyama. 2002. “Male Reproductive Success in a Promiscuous Armoured Catfish *Corydoras Aeneus* (Callichthyidae).” Environmental Biology of Fishes 63 (3): 281–87. https://doi.org/10.1023/A:1014317009892.

Kramer, Annemarie, James L. Van Tassell, and Robert A. Patzner. 2009. “A Comparative Study of Two Goby Shrimp Associations in the Caribbean Sea.” Symbiosis 49 (3): 137–41. https://doi.org/10.1007/s13199-009-0045-7.

LaGory, Kirk E. 1987. “The Influence of Habitat and Group Characteristics on the Alarm and Flight Response of White-Tailed Deer.” Animal Behaviour 35 (1): 20–25. https://doi.org/10.1016/S0003-3472(87)80206-3.

Lambourne, Derek. 1995. Corydoras Catfish: An Aquarist’s Handbook. Blandford, London.

Magurran, Anne E. 1986. “Predator Inspection Behaviour in Minnow Shoals: Differences between Populations and Individuals.” Behavioral Ecology and Sociobiology 19 (4): 267–73. https://doi.org/10.1007/BF00300641.

Magurran, Anne E., and Anthony Higham. 1988. “Information Transfer across Fish Shoals under Predator Threat.” Ethology 78 (2): 153–58. https://doi.org/10.1111/j.1439-0310.1988.tb00226.x.

Magurran, Anne E., and Benoni H. Seghers. 1994. “Predator Inspection Behaviour Covaries with Schooling Tendency Amongst Wild Guppy, Poecilia Reticulata, Populations in Trinidad.” Behaviour 128 (1/2): 121–34.

Major, Peter F. 1978. “Predator-Prey Interactions in Two Schooling Fishes, Caranx Ignobilis and Stolephorus Purpureus.” Animal Behaviour 26 (August): 760–77. https://doi.org/10.1016/0003-3472(78)90142-2.

Neill, S. R. J., and J. M. Cullen. n.d. “Experiments on Whether Schooling by Their Prey Affects the Hunting Behaviour of Cephalopods and Fish Predators.” Journal of Zoology 172 (4): 549–69. https://doi.org/10.1111/j.1469-7998.1974.tb04385.x.

Pitcher, T. J., D. A. Green, and A. E. Magurran. n.d. “Dicing with Death: Predator Inspection Behaviour in Minnow Shoals.” Journal of Fish Biology 28 (4): 439–48. https://doi.org/10.1111/j.1095-8649.1986.tb05181.x.

Preston, J. Lynn. 1978. “Communication Systems and Social Interactions in a Goby-Shrimp Symbiosis.” Animal Behaviour 26 (August): 791–802. https://doi.org/10.1016/0003-3472(78)90144-6.

Radford, A. N. n.d. “Vocal Coordination of Group Movement by Green Woodhoopoes (Phoeniculus Purpureus).” Ethology 110 (1): 11–20. https://doi.org/10.1046/j.1439-0310.2003.00943.x.

Riley, R.J., Gillie, E.R., Johnstone, R.A., Boogert, N.J., Manica, A., 2018. Coping with strangers: how familiarity and active interactions shape group coordination in Corydoras aeneus. bioRxiv 448068. https://doi.org/10.1101/448068

Rutte, C., Taborsky, M., 2007. Generalized Reciprocity in Rats. PLOS Biology 5, e196. https://doi.org/10.1371/journal.pbio.0050196

Sands. 1986. Extracts from The Revision of Surinam Catfish of the Genus Corydoras. Catfishes of the World. Volume One: Supplements (Second Set). Dunure Enterprises, Dunure, Scotland.

Schel, Anne Marijke, Agnès Candiotti, and Klaus Zuberbühler. 2010. “Predator-Deterring Alarm Call Sequences in Guereza Colobus Monkeys Are Meaningful to Conspecifics.” Animal Behaviour 80 (5): 799–808. https://doi.org/10.1016/j.anbehav.2010.07.012.

Seyfarth, R. M., D. L. Cheney, and P. Marler. 1980. “Monkey Responses to Three Different Alarm Calls: Evidence of Predator Classification and Semantic Communication.” Science 210 (4471): 801–3. https://doi.org/10.1126/science.7433999.

Smith, R. Jan F. 1992. “Alarm Signals in Fishes.” Reviews in Fish Biology and Fisheries 2 (1): 33–63. https://doi.org/10.1007/BF00042916.

Sogard, Susan M., and Bori L. Olla. 1997. “The Influence of Hunger and Predation Risk on Group Cohesion in a Pelagic Fish, Walleye Pollock Theragra Chalcogramma.” Environmental Biology of Fishes 50 (4): 405–13. https://doi.org/10.1023/A:1007393307007.

Stensmyr, Marcus C., and Florian Maderspacher. 2012. “Pheromones: Fish Fear Factor.” Current Biology 22 (6): R183–86. https://doi.org/10.1016/j.cub.2012.02.025.

Templeton, Christopher N., Erick Greene, and Kate Davis. 2005. “Allometry of Alarm Calls: Black-Capped Chickadees Encode Information About Predator Size.” Science 308 (5730): 1934–37. https://doi.org/10.1126/science.1108841.

Turchin, P., and P. Kareiva. 1989. “Aggregation in Aphis Varians: An Effective Strategy for Reducing Predation Risk.” Ecology 70 (4): 1008–16. https://doi.org/10.2307/1941369.

Viscido, Steven V., and David S. Wethey. 2002. “Quantitative Analysis of Fiddler Crab Flock Movement: Evidence for ‘Selfish Herd’ Behaviour.” Animal Behaviour 63 (4): 735–41. https://doi.org/10.1006/anbe.2001.1935.

